# rest2vec: Vectorizing the resting-state functional connectome using graph embedding

**DOI:** 10.1101/2020.05.10.085332

**Authors:** Zachery D. Morrissey, Liang Zhan, Olusola Ajilore, Alex D. Leow

**Affiliations:** Graduate Program in Neuroscience, University of Illinois at Chicago; Dept. of Psychiatry, University of Illinois at Chicago; Dept. of Bioengineering, University of Illinois at Chicago; Dept. of Computer Science, University of Illinois at Chicago; Dept. of Electrical and Computer Engineering, University of Pittsburgh

## Abstract

Resting-state functional magnetic resonance imaging (rs-fmri) is widely used in connectomics for studying the functional relationships between regions of the human brain. rs-fmri connectomics, however, has inherent analytical challenges, such as accounting for negative correlations. In addition, functional relationships between brain regions do not necessarily correspond to their anatomical distance, making the intrinsic geometry of the functional connectome less well understood. Recent techniques in natural language processing and machine learning, such as word2vec, have used embedding methods to map high-dimensional data into meaningful vector spaces. Inspired by this approach, we have developed a graph embedding pipeline, rest2vec, for studying the intrinsic geometry of functional connectomes. We demonstrate how rest2vec uses the phase angle spatial embedding (phase) method with dimensionality reduction techniques to embed the functional connectome into lower dimensions. Rest2vec can also be linked to the maximum mean discrepancy (mmd) metric to assign functional modules of the connectome in a continuous manner, improving upon traditional binary classification methods. Together, this allows for studying the functional connectome such that the full range of correlative information is preserved and gives a more informed understanding of the functional organization of the brain.

## 1 Introduction

Neuroimaging data acquired from magnetic resonance imaging (mri) tend to be vast and high-dimensional. In particular, resting-state functional mri (rs-fmri) produces temporal snapshots of the brain’s default activity in the absence of tasks, offering a window into the functional macroscale organization of the brain. As computational tools have become more widely available over the past two decades, researchers have applied graph theory-based models to neuroimaging data to study the network properties of the brain, which has grown into the field of connectomics [1]. In connectomics analyses, the brain can be represented as an *N* × *N* matrix, where the rows and columns are the *N* brain regions of interest (roi), and the elements of the matrix represent some measure of connection between them (e.g., number of fibers, Pearson correlation of blood oxygenation level-dependent (bold) time series). Given this volume of high-dimensional data, however, one quickly runs into the “curse of dimensionality.” Originally coined by Richard Bellman [2], the term refers to the challenge of visualizing and analyzing high-dimensional data. Because the number of points in a Cartesian space grows exponentially with increasing dimensions, high-dimensional spaces become extremely sparse, an effect known as the “empty space phenomenon.” Consequently, this makes understanding the properties of these data more difficult, as metric comparisons become less effective with increasing dimensionality [3].

There are a variety of dimensionality reduction techniques that address this problem. These methods work by embedding a high-dimensional manifold, represented by the discrete points of the data, into a lower dimension (e.g., two or three dimensions), which can then be visualized. This process becomes complicated, however, if the manifold of the underlying data is nonlinear, as is thought to be the case with neuroimaging data [4, 5, 6, 7]. The most well-known example case of a nonlinear manifold is the 3d Swiss roll. Nonlinear dimensionality reduction techniques, such as isometric mapping (isomap), solve the characteristic Swiss roll problem by preserving the intrinsic geometry of nonlinear manifolds (i.e., unrolling the Swiss roll) in lower-dimensional spaces [8, 4].

Negative correlations also remain a challenging factor in rs-fmri connectomics, as they are more difficult to interpret using network models. Simpler models generally either threshold out or apply other transformations to negative correlations, such as taking the absolute value; this process, however, likely removes substantive dynamics of brain connectivity [9]. Although some analyses account for negative correlations, these often introduce additional parameters that must be set to determine their relative contribution [9].

Previously, we introduced probability-associated community estimation (pace) [10] and phase angle spatial embedding (phase) [11] to address these challenges. These methods take inspiration from the Ising model from statistical mechanics, where magnetic ions are designated with in-phase or out-of-phase spin state configurations [12]. We adapted this model to describe the phase relationship between regions of the brain. Here we propose a novel graph embedding pipeline, rest2vec, that uses this phase angle representation with the nonlinear dimensionality reduction method isomap to embed the functional connectome in a lower-dimensional embedding based on its functional relationships. Doing so revealed a spatial mapping of the functional organization of the brain based on its intrinsic geometry when it is not constrained by neuroanatomy.

Additionally, we show this vectorized approach has implications for detecting functional communities by linking rest2vec to the maximum mean discrepancy (mmd) metric. This was originally developed by Gretton et al. [13] as a metric describing the distance between probability distributions. Here, we treated the mmd as a modularity index, similar to *Q*-based maximization methods [14], such that, when maximized, it detects the sets of brain regions with the most dissimilar functional connectivity. By reformulating this connectome modularity problem in a probabilistic sense, we are able to generate community assignment values for each region, as opposed to a binary classification. Together, rest2vec uses nonlinear dimensionality reduction and manifold learning techniques to represent the functional connectome in its intrinsic geometry independent of neuroanatomy to improve our understanding of the macroscale organization of the brain.

## 2 Methods

### 2.1 Dataset description

Two independent and publicly available rs-fmri connectome datasets composed of healthy subjects were used: one from the Functional 1000 (F1000) Connectomes Project [15] with 177 regions of interest (roi) available through the usc Multimodal Connectivity Database (http://umcd.humanconnectomeproject.org/umcd/default/index), and one by Diez et al. [16] with 2514 roi available through the NeuroImaging Tools & Resources Collaboratory (nitrc) (https://www.nitrc.org/projects/biocr_hcatlas/). These are referred to as the “F1000” and “Diez” datasets hereafter. The average difference in age between male (*N* = 426, *M* ± SD = 28.7 ± 12.7) and female (*N* = 560, *M* ± SD = 27.9 ± 12.7) subjects in the F1000 dataset was 0.83 years and was not statistically significant (*t*(984) = 1.025, *p* = 0.306). The Diez dataset has 12 subjects (6 male) with a mean age of 33.5 ± 8.7 years; no individual subject ages were reported. The reader can consult the references for details regarding image acquisition parameters and preprocessing. For computational and network analyses, Python version 3.7.3 scientific computing libraries from the Anaconda distribution were used [17, 18, 19, 20, 21, 22, 23].

### 2.2 rest2vec

The pipeline for rest2vec is shown in Figure 1. Rest2vec aims to create a graph embedding of rs-fmri connectomes by transforming positive and negative edges into *N* -dimensional phase angle vectors that can then be represented in a low-dimensional embedding using nonlinear dimensionality reduction. In brief, we first computed the probability of observing a negative edge between all pairs of regions across all subjects to form the probability matrix **P**^−^. This probability is then used to determine the phase angle Θ_*i,j*_ between regions to create the phase angle spatial embedding (phase) matrix **Θ**. This process embeds the phase relationship between all regions in the connectome in an *N* -dimensional Euclidean space and transforms the values between 0 (fully co-activating) and *π/*2 (fully anti-activating). The intrinsic functional embedding of the connectome was then visualized in two dimensions using the nonlinear dimensionality reduction method isomap [8]. Finally, we use kernel functions to link rest2vec to the maximum mean discpreancy mmd metric [13] to demonstrate how rest2vec can be used to study functional connectome modularity. The representative matrices for each step are displayed in Figure 2.

**Figure 1:**
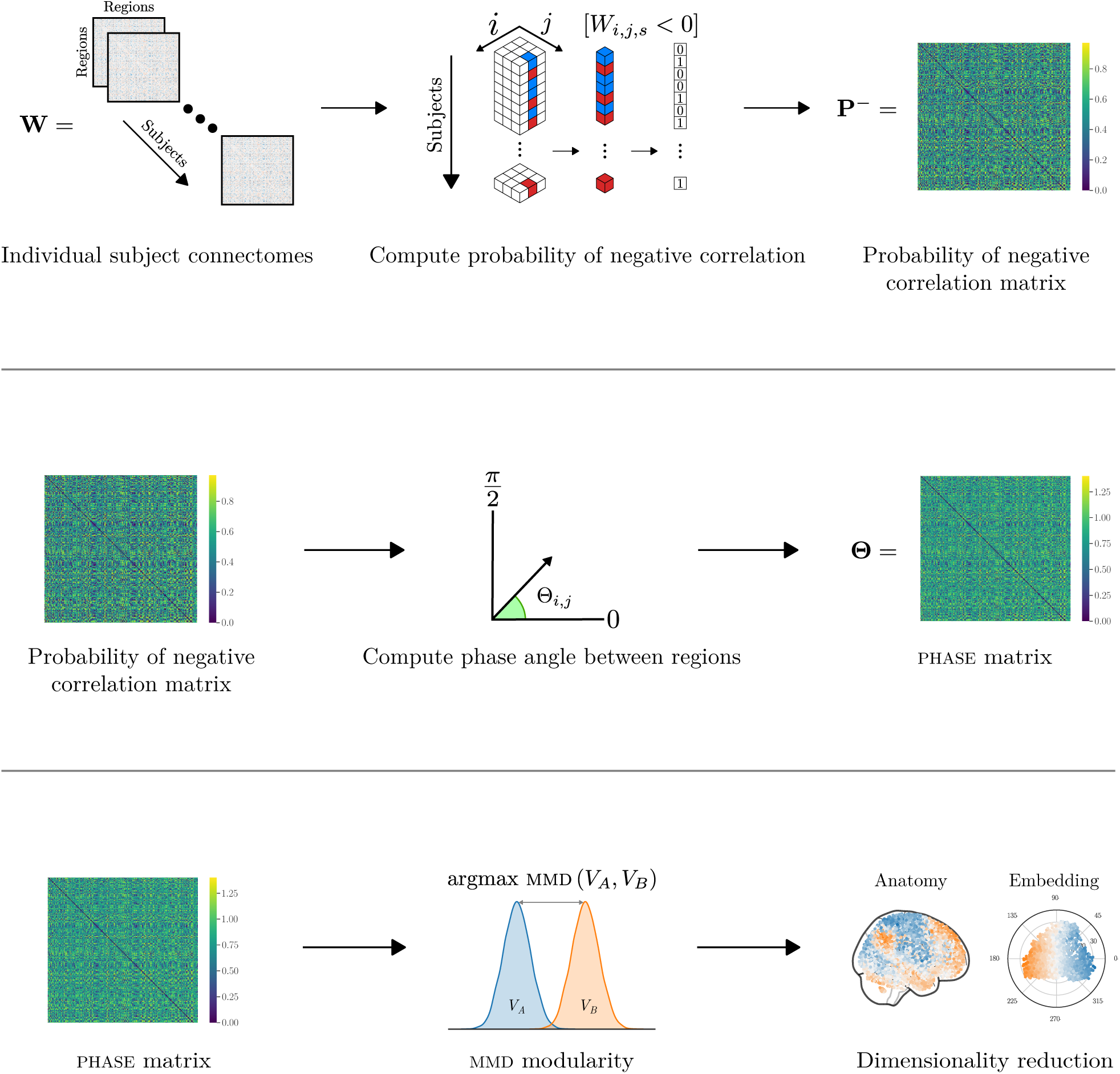
rest2vec processing pipeline. (Top) The frequency of observing a negative edge between regions *i* and *j* across all subjects in the *N* × *N* × *S* array **W** of rs-fmri connectomes is computed to form the probability of negative correlation matrix **P**^−^. (Middle) The phase angle transformation is applied to compute the phase angle spatial embedding (phase) matrix **Θ**. (Bottom) Dimensionality reduction and mmd modularity are used to analyze the properties of the new embedding space, where the functional connectome is represented by its intrinsic geometry.

**Figure 2:**
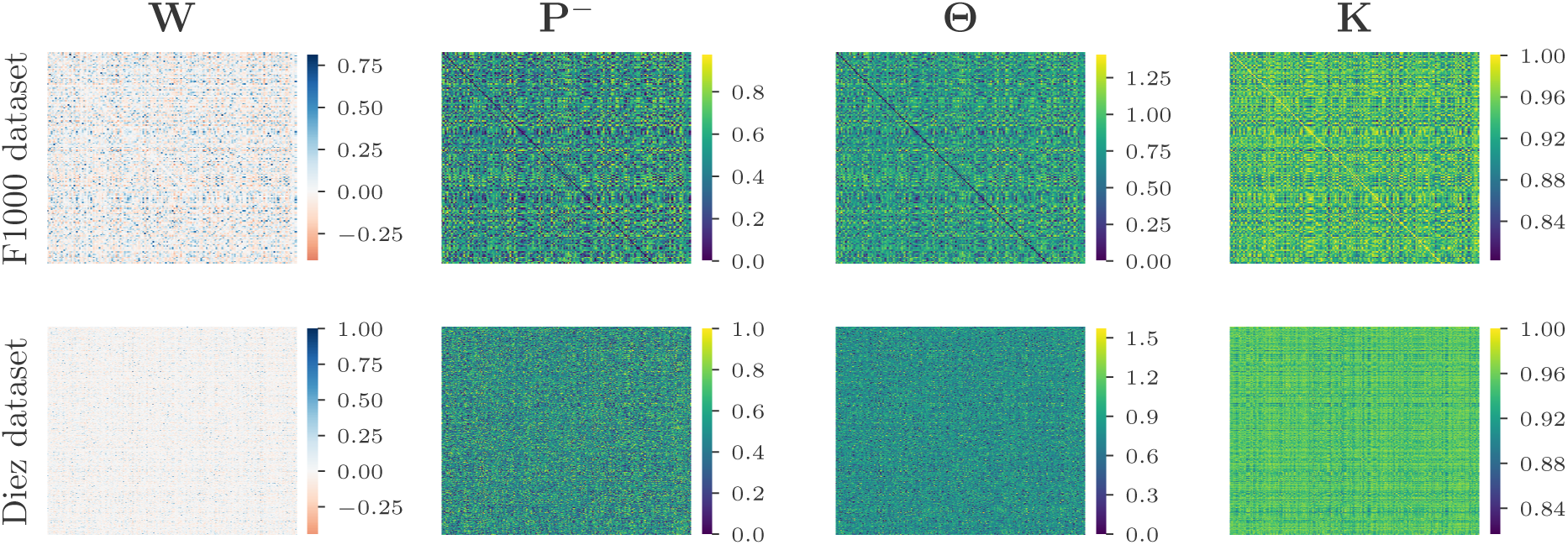
Representative matrices for processing steps of rest2vec pipeline for each dataset. Pearson correlation matrix **W**, negative probability matrix **P**^−^, phase angle spatial embedding (phase) matrix **Θ**, and kernel similarity matrix **K** are displayed.

#### 2.2.1 Phase angle spatial embedding (phase)

A functional connectome derived from rs-fmri is defined as an undirected graph *G*(*V, E*), composed of a set of vertices *V*, i.e., brain regions of interest (roi), and signed, weighted edges *E* describing the measure of connectivity between them based on their bold response time series. Typically, some measure of correlation, e.g., Pearson correlation, between bold time series is used to describe the functional connectivity between roi.

Previously, we introduced probability associated community estimation (pace) [10], and phase angle spatial embedding (phase) [11] for encoding resting-state fMRI connectomes based on the phase relationship between brain regions [11] to account for negative correlations in functional connectomes. We begin by briefly summarizing these procedures in the context of rest2vec.

Let **W** be an *N* × *N* × *S* array (i.e., a tensor) composed of *N* × *N* weighted, signed functional connectomes for *N* regions and *S* subjects. Given some weight of functional coupling between regions *i* and *j* (e.g., Pearson correlation), we define the probability of negative correlation matrix **P**^−^ where each element 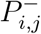 is the probability of observing a negative edge between *i* and *j* defined as 

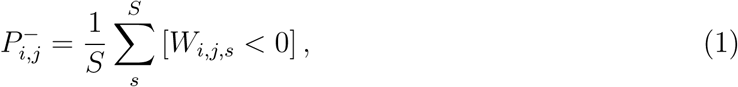

where *W*_*i,j,s*_ is the edge between regions *i* and *j* for the *s*th subject, and the Iverson bracket expression [*W*_*i,j,s*_ < 0] equals 1 if *W*_*i,j,s*_ < 0, and 0 otherwise. Because 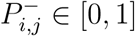, it also follows naturally that

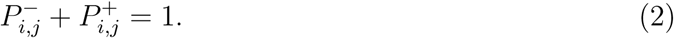

One advantage of this procedure is that the probability measure defined in Equation 1 can be defined by the user for their specific context. By taking advantage of this relationship, we then define the phase angle spatial embedding (phase) matrix **Θ**, where the phase angle Θ_*i,j*_ between regions *i* and *j* is defined as

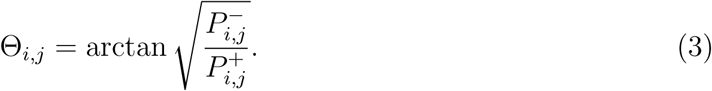

Thus Θ_*i,j*_ ∈ [0, *π/*2], where 0 represents a fully in-phase (co-activating) relationship and *π/*2 represents a fully out-of-phase (anti-activating) relationship. Each column of **Θ** is a vector embedding each region in an *N* -dimensional Euclidean space such that **Θ**_:,*i*_ = [Θ_*i*,1_ Θ_*i*,2_ … Θ_*i,N*_]^T^ ∈ [0, *π/*2]

#### 2.2.2 Relation of PhASE to the maximum mean discrepancy

Here we describe how phase can be linked to the maximum mean discrepancy (mmd) developed by Gretton et al. [13] to address the connectome modularity problem. Following the formulation defined in [13], consider the random variables *x* and *y* defined on a metric space 𝒳 equipped with the metric *d*, with the corresponding Borel probabilities *p* and *q* (i.e., *x* ∼ *p* and *y* ∼ *q*). Given observations *X* := {*x*_1_, …, *x*_*m*_} and *Y* := {*y*_1_, …, *y*_*n*_} drawn from the probability distributions *p* and *q, p* = *q* if and only if **E**_*x*_[*f* (*x*)] = **E**_*y*_[*f* (*y*)] ∀*f* ∈ *C*(𝒳), where *C*(𝒳) is the space of bounded continuous functions on 𝒳. Next, given a class of functions ℱ such that *f* : 𝒳 → ℝ, the maximum mean discrepancy (mmd) between *p* and *q* with respect to ℱ is defined as

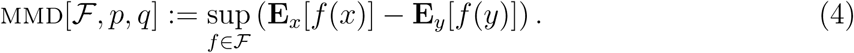

This can be empirically estimated given *X* and *Y* as

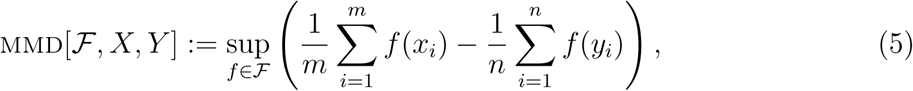

where *m* is equal to the number of observations in *X* and *n* is equal to the number of observations in *Y*.

To apply these definitions in the context of connectomics, we use the same definitions of *x, y, p, q, X*, and *Y* defined above to assign each region to one of the two distributions *p* or *q*.

Under the working assumption that the distributions of functional modules in the connectome are far apart (i.e., their within-module connections are greater than their between-module connections [24]), we thus seek to discover the arrangement of regions such that the mmd between them is maximized.

Using a reproducible kernel Hilbert space (rkhs), the squared form of Equation 5 can be evaluated using kernel functions as

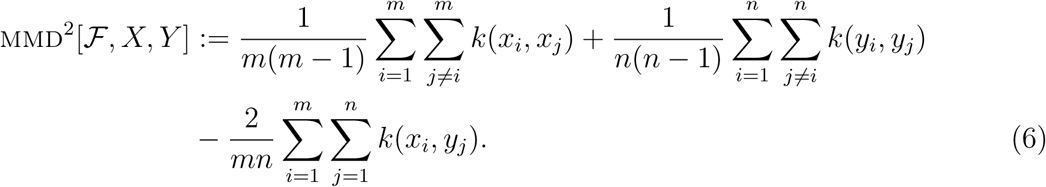

From Equation 6, kernel functions can be used, in our case, to compute the kernel matrix **K** where the similarity *K*_*i,j*_ between regions *i* and *j*, in the case of the radial basis function (rbf) kernel *k*_RBF_, is given by

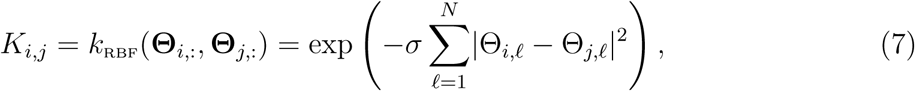

for phase angle Θ between regions *i* and *j* in reference to all other regions indexed by *f*, for *N* regions, using the scaling factor *σ*.

Similarly, we let the cosine kernel *k*_cos_ evaluating the similarity between regions *i* and *j* be defined as

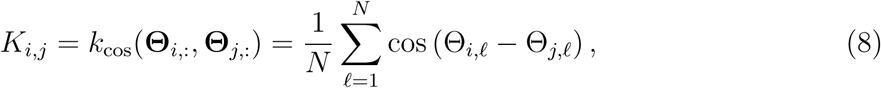

using the same variable definitions as rbf kernel. Because the rbf kernel has an additional parameter, and the cosine kernel has a geometric relation to angles, the cosine kernel is used here; the Taylor expansion of both these kernels can be shown to have similar leading terms.

#### 2.2.3 Using maximum mean discrepancy to address the connectome modularity problem

Following the kernel definitions above, and the equation as described by [13] (with a modified notation for our purposes), let the maximum mean discrepancy (mmd) between two modules *V*_*A*_ and *V*_*B*_ be defined as

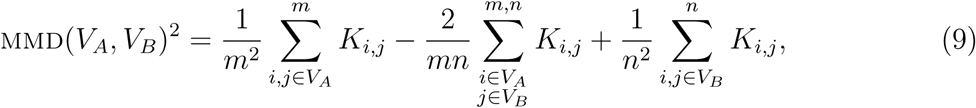

where |*V*_*A*_| = *m*, |*V*_*B*_| = *n*, |*V* | = *m* + *n* = *N, V*_*A*_ ∪*V*_*B*_ = *V, V*_*A*_ ∩*V*_*B*_ = Ø, and *i* is allowed to equal *j*.

We seek to find a partition between *V*_*A*_ and *V*_*B*_ such that Equation 9 is maximized. First, we can rewrite mmd(*V*_*A*_, *V*_*B*_)^2^ = **y**^T^**Ky** for **y** ∈ ℝ^*N* ×1^ and **K** ∈ ℝ^*N*×*N*^, where

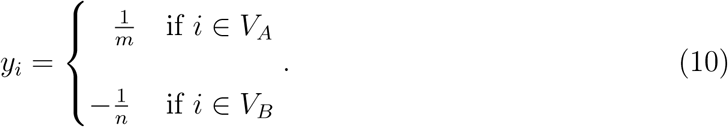

Thus we define the optimal partition Modularity(*V*) into modules *V*_*A*_ and *V*_*B*_ as

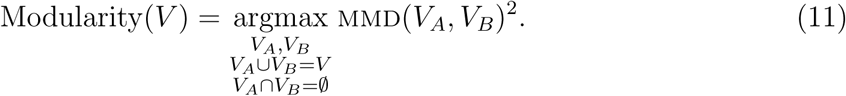

This maximization problem can be approximated in a simplified way by relaxing Equation 11 to a Rayleigh quotient maximization problem. Letting **y** be defined as above, where

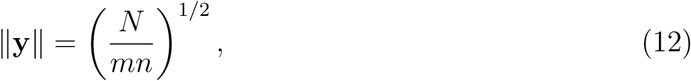

we perform change of variables to the unit length vector **v**, where

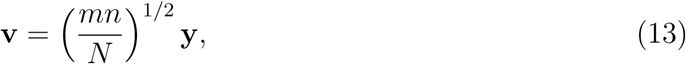

and ‖**v**‖^2^ = 1, **v**^T^**1** = 0, where **1** = [1 … 1]^T^, **1** ∈ ℝ^*N* ×1^. Then we can rewrite Equation 11 in terms of **v** to define the partition that maximizes mmd(*V*_*A*_, *V*_*B*_)^2^ as

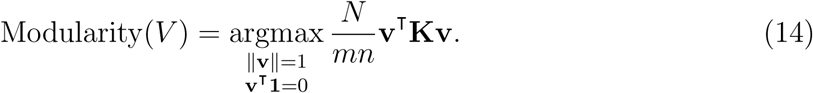

To compute mmd(*V*_*A*_, *V*_*B*_)^2^ in Equation 14 requires *a priori* knowledge of *m* and *n*. Assuming that *N* is large and that the two communities *V*_*A*_ and *V*_*B*_ are approximately the same size such that |*m* − *n*| ∈ *o*(*N*), the normalization factor in Equation 14 can be simplified to

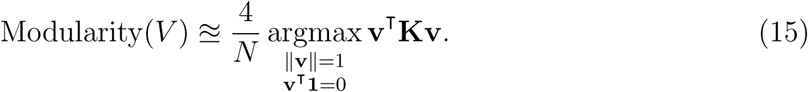

Finally we relax the constraints of **v** from 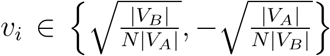 taking only two values to taking any real values such that **v**^∗^ ∈ ℝ^*N*^. These relaxed constraints allow us to conveniently reframe Equation 14 as a Rayleigh quotient maximization problem. We account for arbitrary origin for the Rayleigh quotient maximization by centering the kernel similarity matrix **K** to 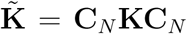, where the centering matrix 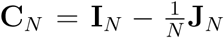, **C**_*N*_ ∈ ℝ^*N*×*N*^, **I**_*N*_ ∈ ℝ^*N*×*N*^ is the identity matrix, and **J**_*N*_ ∈ ℝ^*N*×*N*^ is the ones matrix (i.e., **11**^T^).

Rather than finding mmd(*V*_*A*_, *V*_*B*_)^2^ as a function of the partition, we approximate the optimal partition Modularity(*V*) by finding the vector **v**^∗^ that maximizes the Rayliegh quotient such that

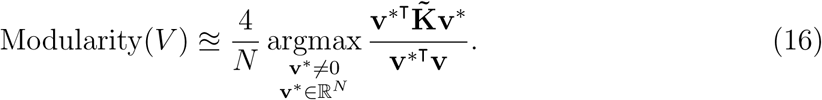

We can then compute the mapping vector **v**^∗^ that maximizes the Rayleigh quotient by computing the eigenvector **q** of 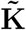 corresponding to the largest eigenvalue λ_max_ of 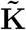. Similar to the Fiedler vector in spectral clustering methods [25], the elements of **q** assign both community affiliation based on its sign (+ or −) as well as magnitude. Further, **q** can be binarized to determine discrete community labels for each region as

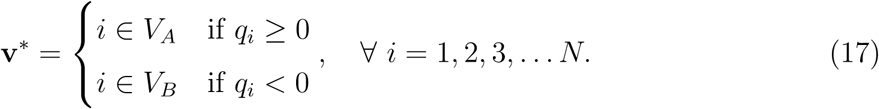

#### 2.2.4 Nonlinear dimensionality reduction

Isomap [8] was used to reduce the phase matrix **Θ** ∈ ℝ^*N*×*N*^ to a *d*-dimensional embedding **Y** ∈ ℝ^*N*×*d*^, where *d* < *N*. Isomap is advantageous for this procedure as it is a nonlinear technique, using methods such as Dijkstra’s algorithm [26] to compute the geodesic distances between vertices in high-dimensional space. By doing so, isomap addresses the Swiss roll problem faced by traditional linear methods such as pca and mds [8]. In our case, we used *k* = 12 and *k* = 50 nearest neighbors, for the F1000 and Diez datasets, respectively, to reduce to three dimensions using the Isomap implementation in the Scikit-learn version 0.21.3 library [22]. Because the isomap procedure centers data about the origin, and by Equation 3 the phase angle between perfectly in-phase regions is zero, we analyzed each region’s Euclidean distance to the origin in this space to observe how the phase relationship between regions is preserved with respect to its low-dimensional embedding. After generating the isomap embedding, the distance *D*_*i*_ to the origin of the isomap space [0 … 0] ∈ ℝ^1×*d*^ for the *i*th region was calculated using the Euclidean distance

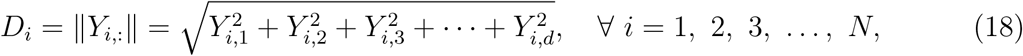

where *N* is equal to the number of regions. Because of its natural representation for distance to the origin, the first two dimensions were transformed to polar coordinates of radius *r* and angle *θ* using the polar transformation

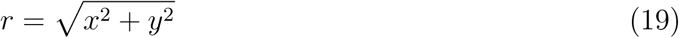

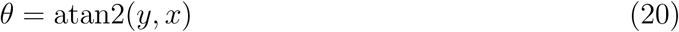

to visualize the functional embedding space.

### 2.3 Analyses

#### 2.3.1 *k*-means clustering

*k*-means clustering was used to formally classify clusters for regions (such as the precuneus) that had heterogeneous mappings in the isomap embedding. The *k*-means clustering algorithm was performed using the Scikit-learn implementation [22] for *k* = 2 clusters in the isomap embedding. The same seed value was used to ensure reproducible results.

To determine how affiliated other (non-precuneus) regions were to either of the two clusters, regions were first assigned to the precuneus cluster they were closest to in the isomap embedding. A diverging cluster affiliation scale was computed based on the Euclidean distance of each region to its precuneus cluster’s centroid in the isomap embedding, which we termed “intrinsic functional distance,” such that regions with more positive or negative values were closer to the centroid of their respective precuneus cluster. The cluster affiliation *a*_*i*_ was defined as

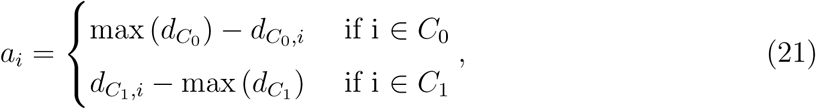

where *d* is the intrinsic functional distance from region *i* to the centroid of cluster *C*.

#### 2.3.2 Statistics

The StatsModels library version 0.10.1 for Python [27] was used for statistical analyses. Student’s independent *t*-test was used to test if there were any differences in age between male and female subjects for the F1000 dataset. The ordinary least squares (ols) method was used to fit the parameters for the linear regression between isomap distance to origin and phase angle.

#### 2.3.3 Visualization

Graphics were drawn using the Matplotlib version 3.1.1 [20] and Seaborn version 0.9.0 [21] libraries using Python version 3.7.3 from the Anaconda distribution [17]. Glass brain figures were visualized using the `plot_connectome` function from the Nilearn version 0.6.2 library [28]. Inkscape version 0.92 was used for final arrangement of some figures [29].

Brain surface plots were created by representing the *N* × 4 array, consisting of the mni (*x, y, z*)-coordinates for all *N* regions, and the *N*×1 vector containing the data value associated with each region, as a 3d volume. For brain distance maps, the intrinsic functional distance vector was made by computing the Euclidean distance between the mean (*x, y*)-coordinates of the anatomical region in the isomap embedding and all other regions. For regions that had heterogeneous mapping (i.e., multiple clusters) in the isomap space, *k*-means clustering was performed to calculate cluster affiliations for each region as described in §2.3.1.

The 3d volume containing the original data was then interpolated using a linear grid interpolation and registered to the mni template volume with 12 degrees of freedom using the `FLIRT` tool in the fsl [30] interface from the Nipype version 1.3.0-rc1 library [31]. The interpolated 3d volume was mapped to the Freesurfer pial surface template using the `vol_to_surf` function from the Nilearn library. The surface data was then visualized using the `plot_surf_stat_map` function from the Nilearn library.

#### 2.3.4 Code

All code used to produce the results and figures is available online via GitHub (https://github.com/zmorrissey) and our laboratory website (http://brain.uic.edu/).

## 3 Results

### 3.1 Distance in the lower-dimensional embedding preserves phase angle relationships

After applying the rest2vec pipeline to the F1000 and Diez datasets, we sought to assess how a region’s lower-dimensional isomap embedding related to its phase vector. From Equation 3, lower values of Θ_*i,j*_ indicate a more in-phase relationship between regions. Thus we hypothesized that more in-phase regions would be embedded closer to the origin of the isomap space, whereas more out-of-phase regions would be embedded further from the origin. The 2-norm of each *N* -dimensional vector of the phase matrix ‖Θ_*i*,:_‖ was used as a summary measure of each region’s overall phase value. For each dataset, there was a statistically significant positive correlation between each region’s ‖**Θ**_*i*,:_‖ and its distance from the origin of the 3d isomap embedding (F1000 dataset: *F* (1, 175) = 200.7, *R*^2^ = 0.534, *r* = 0.731, *p* < 0.0001; Diez dataset: *F* (1, 2512) = 533.4, *R*^2^ = 0.175, *r* = 0.418, *p* < 0.0001) (Figure 3, left). This pattern can be seen when the rows and columns of the phase matrix are sorted by ascending ‖**Θ**_*i*,:_‖ values, in particular for the coarser parcellation from the F1000 dataset (Figure 3, right). Together this suggests that regions mapped closer to the origin were more in-phase with other regions, whereas more out-of-phase regions were mapped further from the origin.

**Figure 3:**
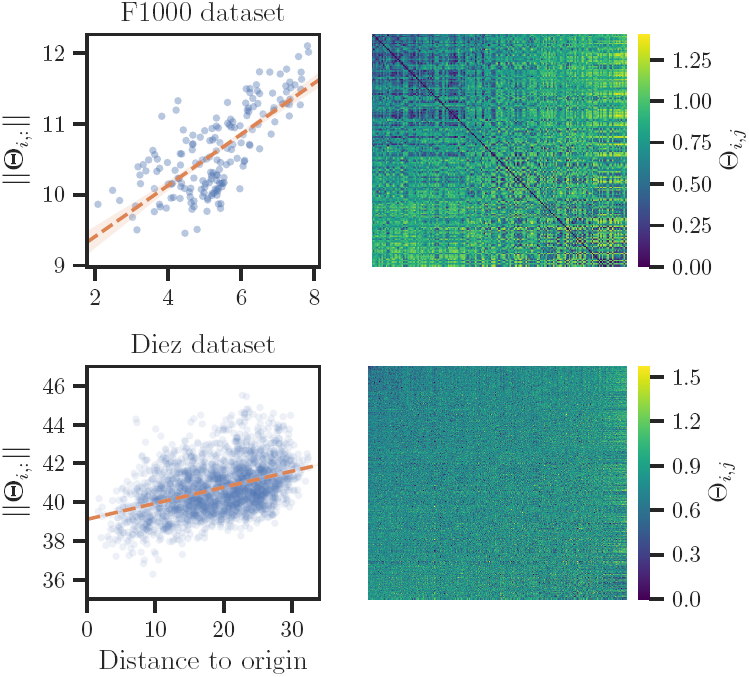
Relationship between phase angle and isomap embedding distance. (Left) Correlation between the 2-norm of each region’s phase angle vector ‖**Θ**_*i*,:_‖ and its distance to the origin of the 3d isomap embedding. Dashed orange line represents the best fit of the linear model. Shaded region around line represents the 95% confidence interval of the model. F1000 dataset: *r* = 0.731; Diez dataset: *r* = 0.418. (Right) The phase matrix **Θ** with its rows and columns sorted in ascending order by ‖**Θ**_*i*,:_‖ (i.e., lowest values correspond to upper left, highest values to lower right).

To examine this relationship further, we faceted the anatomical and functional embeddings by anatomical lobe affiliation ranked by ascending distance to the origin (Figure 4). Notably, the brainstem displayed the most centrally-embedded regions (median distance = 6.9), followed by (in ascending order): sub-lobar, limbic lobe, temporal lobe, frontal lobe, cerebellum, parietal lobe, and occipital lobe regions. At the other extreme, the occipital lobe displayed the most distant and densely clustered representation in the embedding space (median distance = 24). Examination of the phase angle vectors for occipital lobe regions revealed highly in-phase relationships within the occipital lobe, while regions outside the occipital lobe were mostly out-of-phase (SI Figure 15). Since the occipital lobe and large portions of the parietal lobe (e.g., motor cortices), and cerebellum are mapped further in the periphery, this suggests that regions involved in primary sensory processing are mapped further in the periphery, while regions such as the brainstem, thalamus, and heteromodal areas have more in-phase relationships and are mapped closer to the origin.

**Figure 4:**
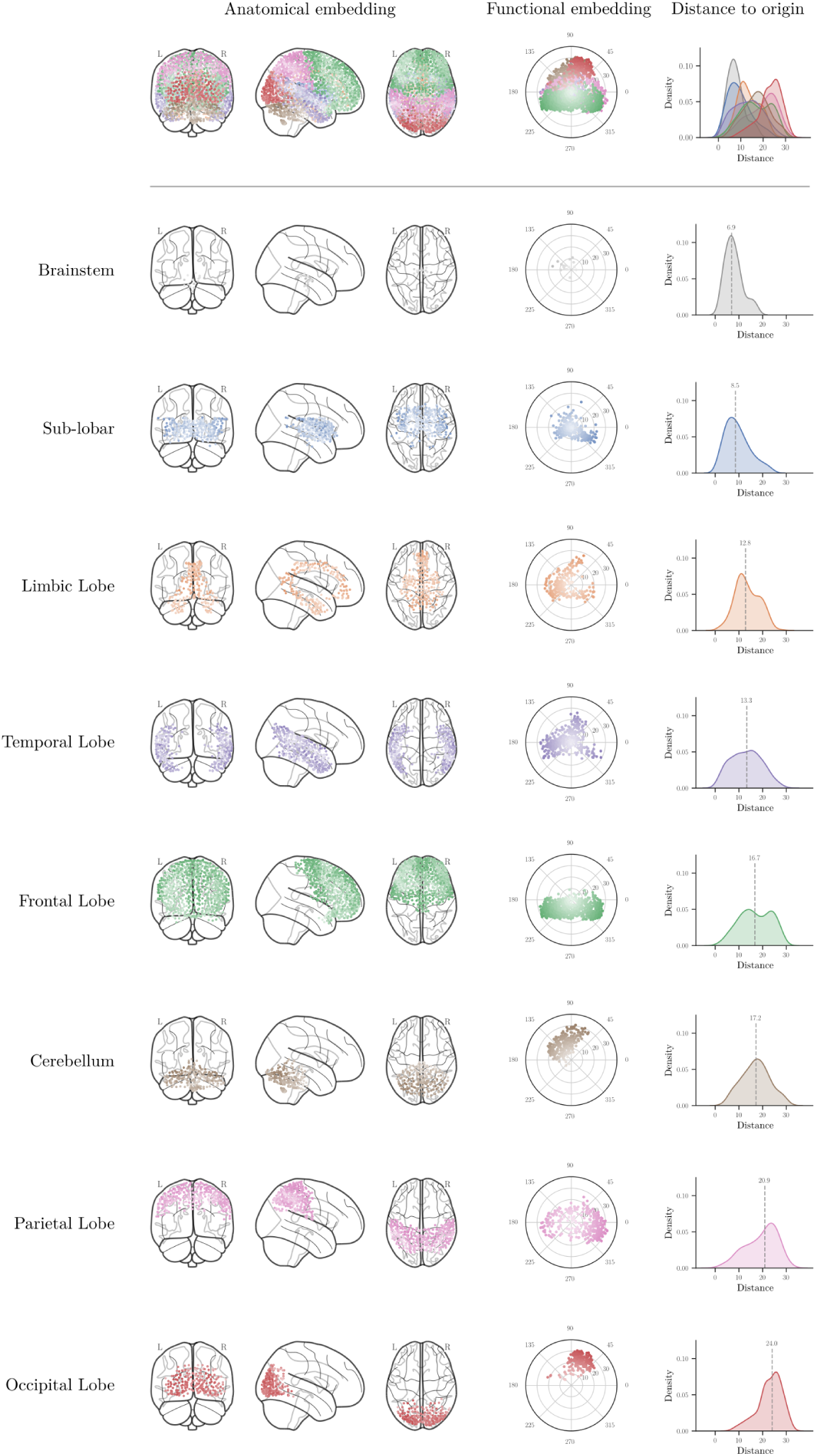
Anatomical and functional embedding of the Diez dataset faceted by anatomical lobe affiliation and ranked by ascending distance to origin. (Top) Merged representations of all 2514 regions in the anatomical embedding (columns 1-3), functional embedding (column 4) and kernel density estimate of distance to origin for all regions within each lobe (column 5). (Bottom) Facet of data for each anatomical lobe. Rows are arranged from top to bottom in ascending order of median distance from the origin from top to bottom. Color indicates lobe affiliation. Higher saturation indicates increasing distance from the origin. Dashed gray lines in kernel density estimate plots indicate the median distance.

### 3.2 Intrinsic functional distance can detect biologically-relevant connectivity gradients

Given that the distance to the origin of the isomap embedding preserved phase coupling characteristics across anatomical regions, we next asked if the intrinsic functional distance between regions in this space could reveal biologically-relevant connectivity patterns. When the intrinsic functional distance to the occipital lobe is mapped as a color gradient on the brain surface, the dorsal and ventral visual streams [32, 33, 34] become apparent (Figure 5), consistent with the hypothesis that distance in this embedding space preserves functionally relevant information. In contrast, the hippocampus also has a relatively homogeneous cluster in the isomap embedding, but has a much more distributed surface map gradient to regions of the default mode network (dmn), such as the precuneus, prefrontal cortex, thalamus, and inferior parietal lobule (Figure 6).

**Figure 5:**
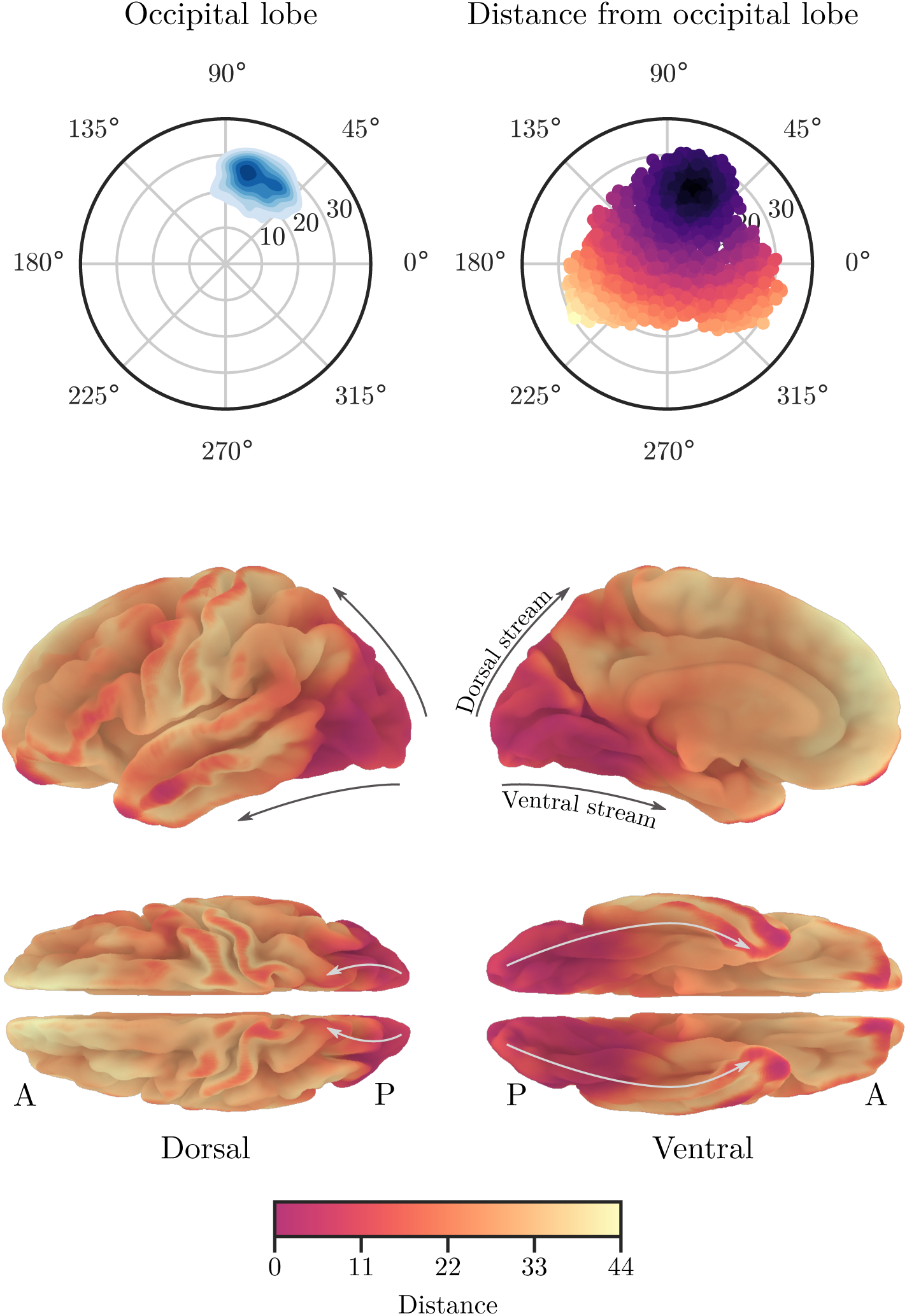
Occipital lobe intrinsic functional distance mapping. (Top left) Kernel density estimate plot of the occipital lobe regions in the isomap embedding. (Top right) Intrinsic functional distance to the occipital lobe for all regions. Darker color indicates the region is closer to the mean occipital lobe coordinate. (Bottom) Intrinsic functional distance to the occipital lobe projected onto the Freesurfer pial surface template. Arrows indicate the dorsal and ventral visual streams. A: anterior. P: posterior.

**Figure 6:**
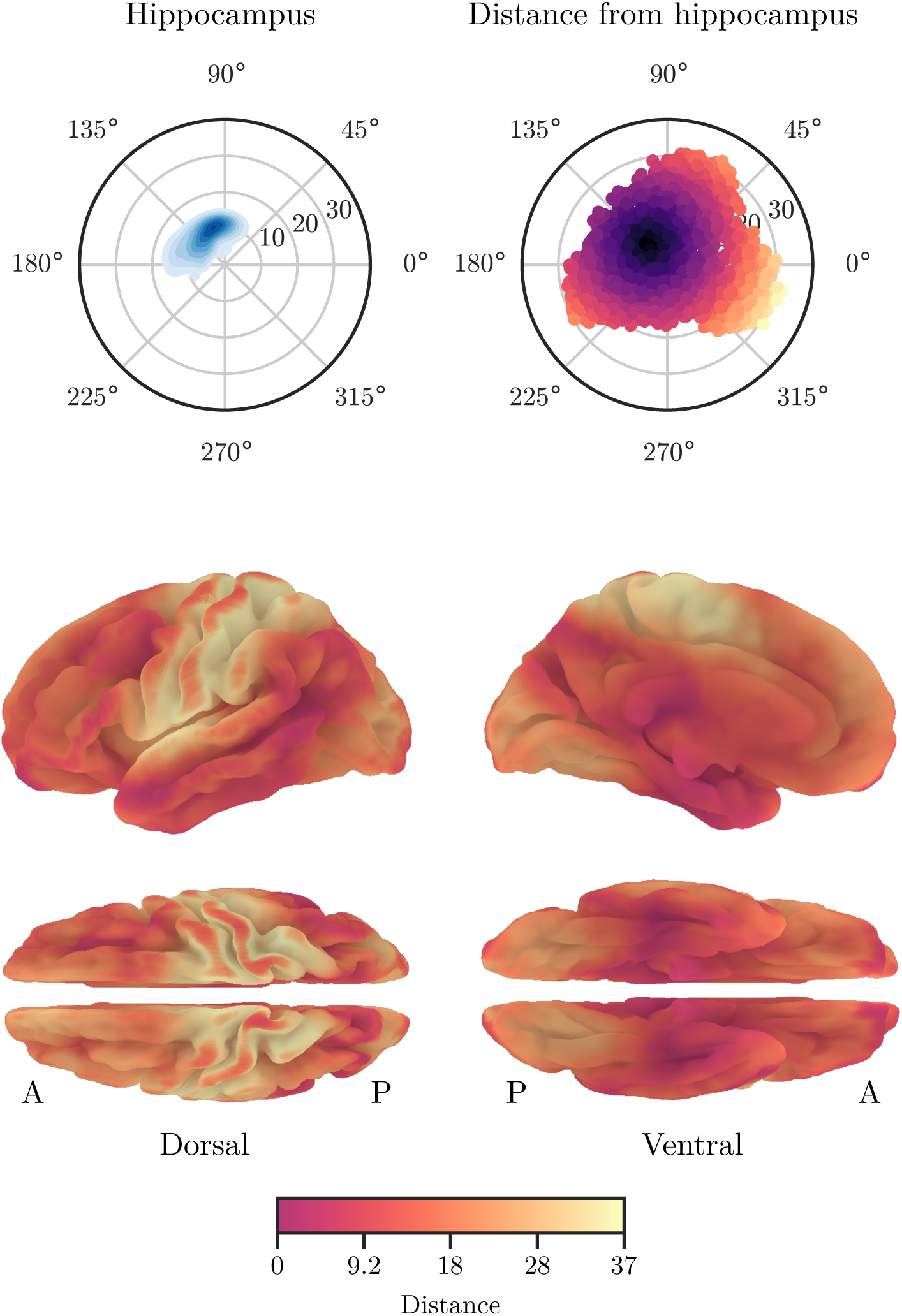
Hippocampus intrinsic functional distance mapping. (Top left) Kernel density estimate plot of the hippocampus regions in the isomap embedding. (Top right) Intrinsic functional distance to the hippocampus for all regions. Darker color indicates the region is closer to the mean hippocampus coordinate. (Bottom) Intrinsic functional distance to the hippocampus projected onto the Freesurfer pial surface template. A: anterior. P: posterior.

While certain anatomical regions showed a relatively homogeneous clustering in the isomap embedding, such as the occipital lobe, others showed heterogeneous clustering patterns. Thus we hypothesized that rest2vec could be used to identify functional subnetworks within individual regions based on their clustering within the isomap embedding. As a test case, we examined the isomap embedding pattern for the precuneus, which is known to participate in different networks across its dorsal-anterior/ventral-posterior axes [35, 36]. The bivariate kernel density estimate plot of the precuneus roi in the Diez dataset appeared to indicate two predominant clusters, which were formally assigned using *k*-means clustering (Figure 7, top). A larger cluster was made that included all other regions in the Diez dataset by assigning regions to the precuneus cluster they were closer to. We then measured the intrinsic functional distance between each region to its precuneus cluster centroid to assign an affiliation value to each region (Figure 7, top right). The brain surface map projection of these data demarcated these two cluster centroids into the dorsal-anterior precuneus and the ventral-posterior precuneus (Figure 7, bottom). The dorsal-anterior cluster of the precuneus was most strongly affiliated with the occipital and superior parietal regions, as well as the paracentral lobule, middle and superior temporal cortices, and thalamus (Figure 7, middle). The ventral-posterior cluster of the precuneus was most strongly affiliated with the hippocampus, cuneus, cerebellum, parahippocampal cortex, posterior cingulate cortex, calcarine cortex, amygdala, and superior occipital cortices. These results suggest that rest2vec can identify distinct functional networks within individual regions.

**Figure 7:**
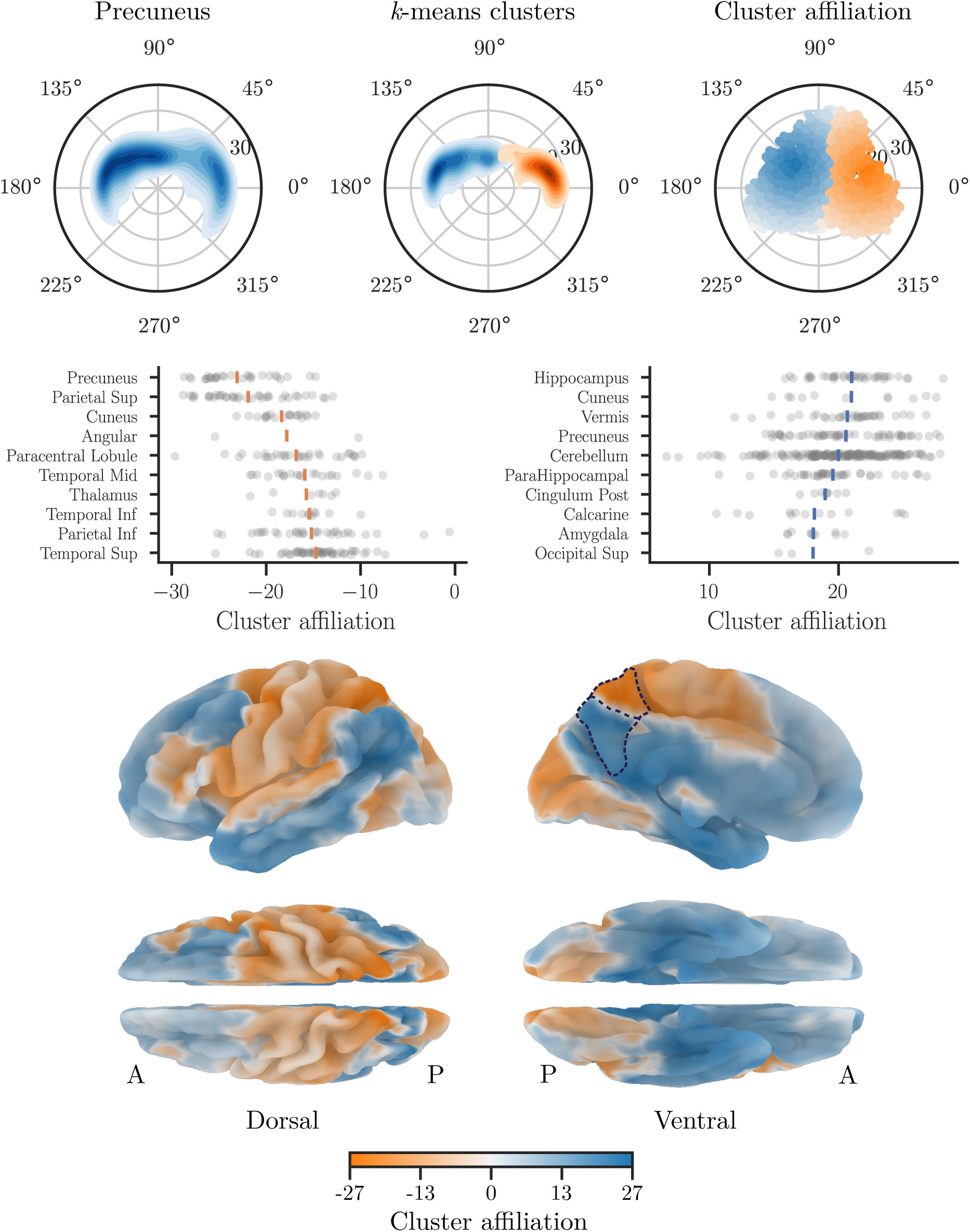
Identifying subnetwork clusters within the precuneus using rest2vec. (Top, left) Kernel density estimate of the precuneus in the isomap embedding. (Top, middle) *k*-means clustering results are indicated in blue and orange. (Top, right) Cluster affiliations for all other regions based on their minimum intrinsic functional distance to their precuneus cluster centroids. Darker color indicates that region is closer to the centroid of its cluster. (Middle) Strip plot of the ten regions with the greatest mean affiliation for each cluster. Points represent individual roi. Vertical bars indicate the mean. (Bottom) Brain surface map of cluster affiliations for the precuneus. The precuneus is outlined by a dashed line in the medial view. A: anterior. P: posterior.

### 3.3 Maximizing maximum mean discrepancy partitions the connectome into putative task-positive and task-negative networks

Since rest2vec could identify functionally relevant connectivity gradients within anatomical lobes, we next asked if rest2vec could be used to partition rs-fmri connectomes into functional modules. To address this, we used the maximum mean discrepancy (mmd) metric developed by [13] to partition the set of connectome regions *V* into two distributions of regions *V*_*A*_ and *V*_*B*_ such that the mmd between them was maximized. A cosine kernel (Equation 8) was used to compute the centered kernel similarity matrix 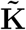 between all pairwise regions of **Θ** (cf. Figure 2). Similar to spectral clustering methods, we approximated the maximum mmd by reformulating the mmd to a Rayleigh quotient maximization problem (§2.2.3), where the eigenvector **q** corresponding to the maximum eigenvalue was extracted to yield the community assignment vector. (Ranking of the eigenvectors of 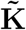 showed the first three eigenvalues account for most of the variance of the data, with the first being most dominant (Figure 8).)

**Figure 8:**
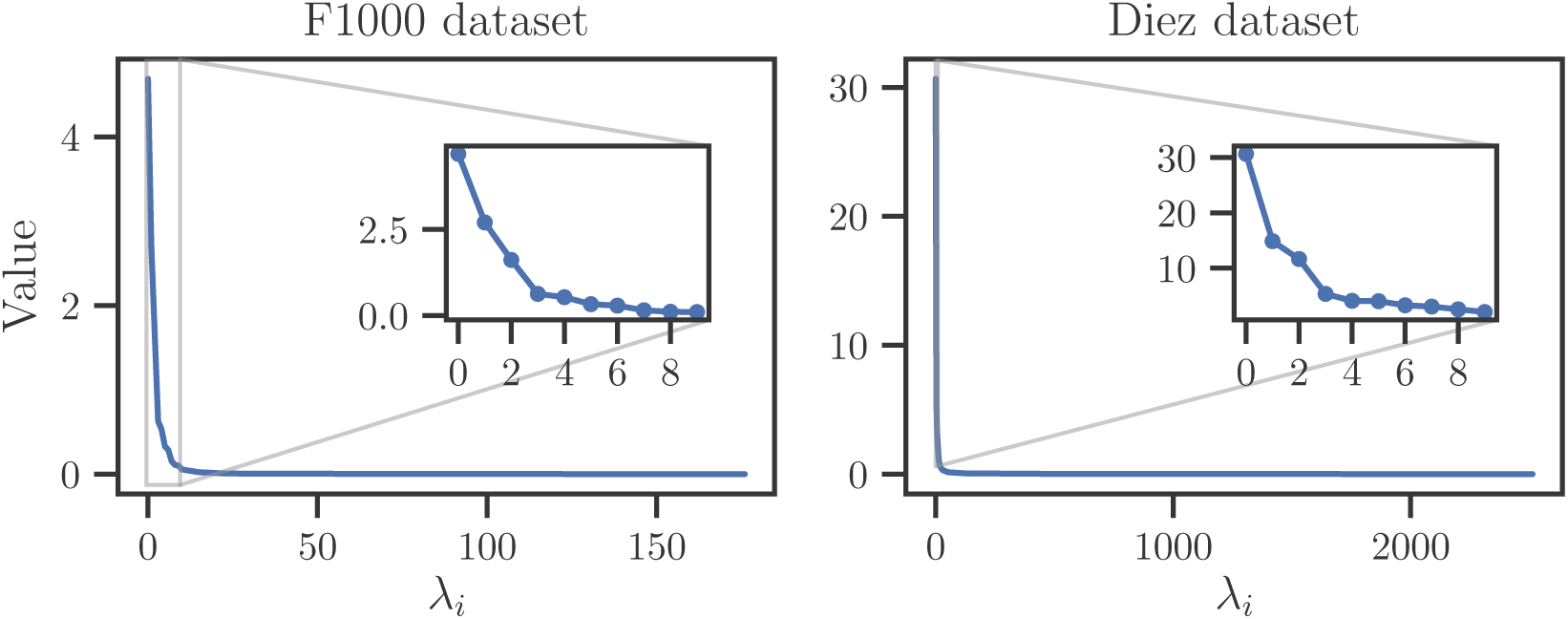
Eigenvalues of 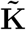 for each dataset. Insets depict the first ten eigenvalues.

To set the partition, the index of regions corresponding to *q*_*i*_ ≥ 0 were assigned to *V*_*A*_, and the index of regions corresponding to *q*_*i*_ < 0 were assigned to *V*_*B*_. This approach was validated by iteratively evaluating the mmd across 50 threshold values of **q** (Figure 9). The results suggest that the mmd is maximized when the partition yields communities of approximately equal size, which occurs for both datasets when the partition threshold for *q*_*i*_ ≈ 0. Together this suggests that the Rayleigh quotient maximization approximation is able to achieve an accurate approximation of the global maximum mmd.

**Figure 9:**
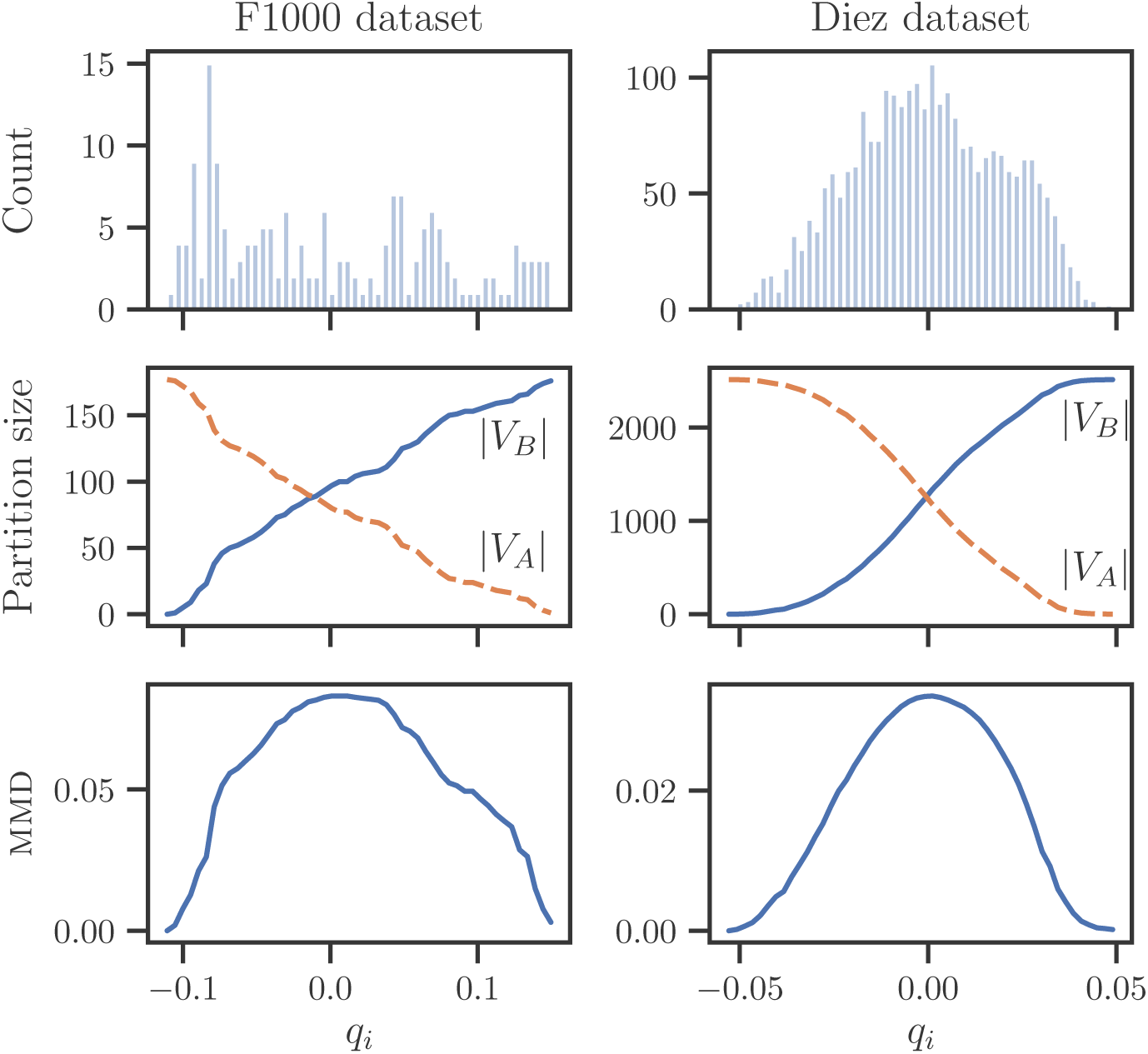
Evaluation of the maximum mean discrepancy (mmd) across a range of threshold values of **q** for each dataset. (Top) Histogram of the eigenvector **q** corresponding to the maximum eigenvalue of the centered kernel matrix 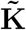. (Middle) Size of each partition as a function of threshold value. (Bottom) mmd as a function of threshold value.

Similar to the previous analysis in Figure 4, we visualized the anatomical and functional embedding by community affiliation to see how each lobe participates in the two communities predicted by maximizing the mmd. We observed a symmetrical partition between the two communities when viewed in the functional embedding space (Figure 10). Additionally, when the magnitude and sign of *q*_*i*_ are mapped to a diverging colormap in the isomap space, it was observed that regions closer to the vertical axis appeared more neutral, whereas regions further from the vertical axis were polarized into either community, suggesting these regions are more strongly mapped into that community. When further examining the regions that are affiliated with each community, we observed that the partition demarcated into the putative task-positive network (tpn) and task-negative network (tnn, also called default mode network (dmn)). This relationship can be seen when the mmd eigenvector gradient is used to sort the **Θ** matrix for each datasets, where the resulting grid communities show out-of-phase relationships with the other community (Figure 11). This can also be observed anatomically when the eigenvector gradient is mapped to the brain surface (Figure 12).

**Figure 10:**
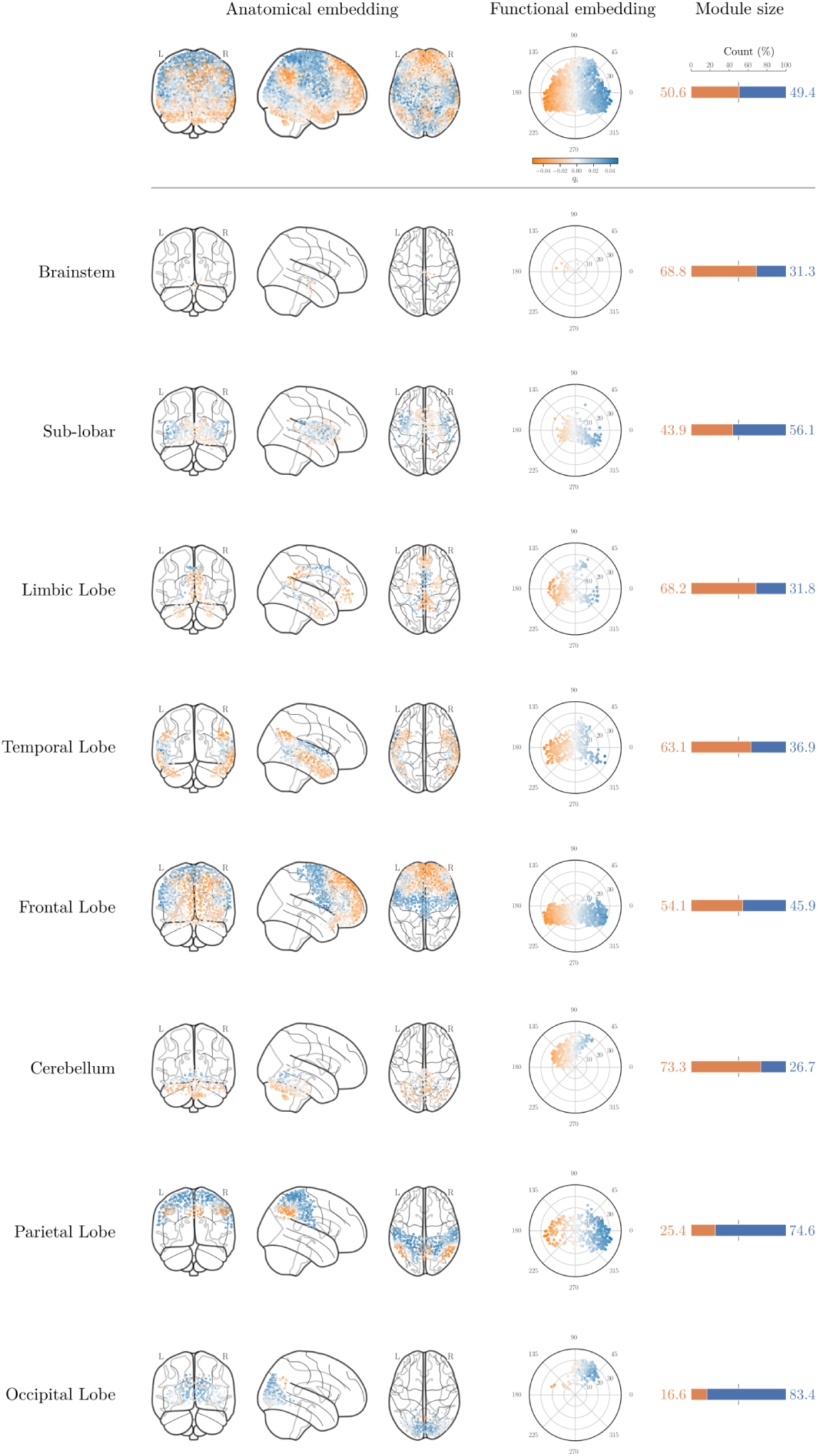
Anatomical and functional embedding of the Diez dataset faceted by anatomical lobe affiliation with predicted community partitions. (Top) Merged representations of all 2514 regions in anatomical embedding (columns 1-3), functional embedding (column 4), and the percentage of regions within each community for each lobe (column 5). (Bottom) Facet of data for each anatomical lobe. Rows are arranged from top to bottom in ascending order of median distance from the origin. Color indicates community affiliation. Vertical gray reference line for each stacked barplot indicates 50%.

**Figure 11:**
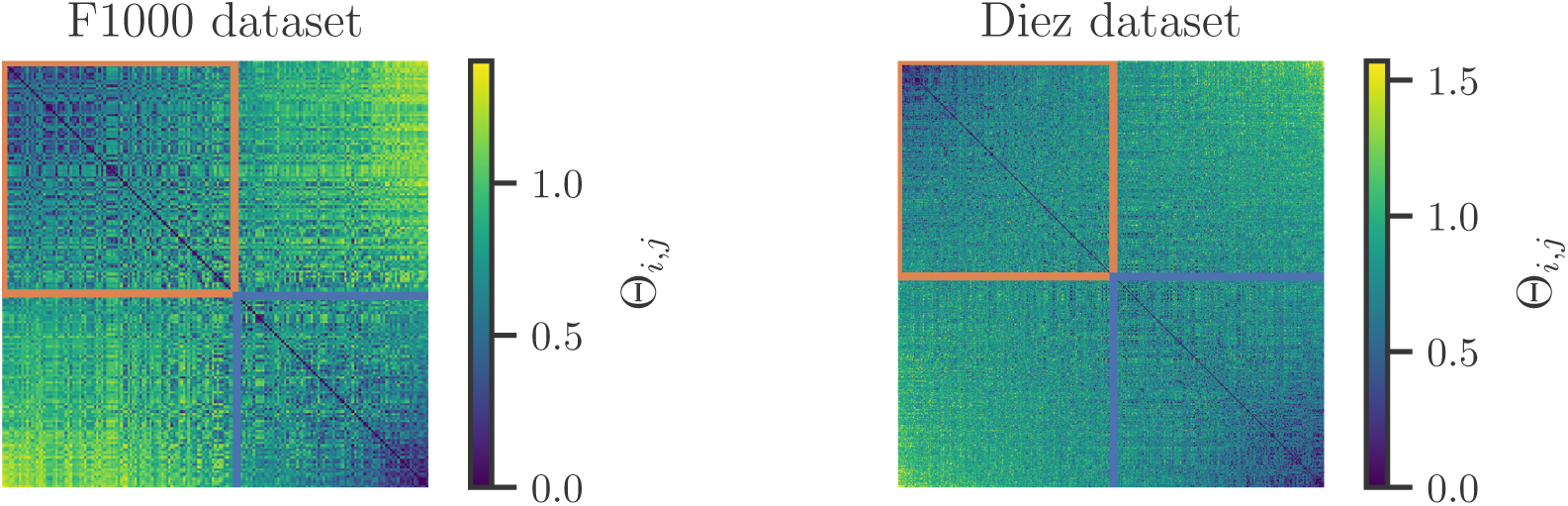
Community partition gradient given by the top eigenvector **q** of the kernel similarity matrix 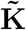. The phase matrix **Θ** for the F1000 and Diez datasets are shown with rows and columns sorted (in ascending order) by the elements of **q**. Grid lines indicate the community partition (upper left and lower right).

**Figure 12:**
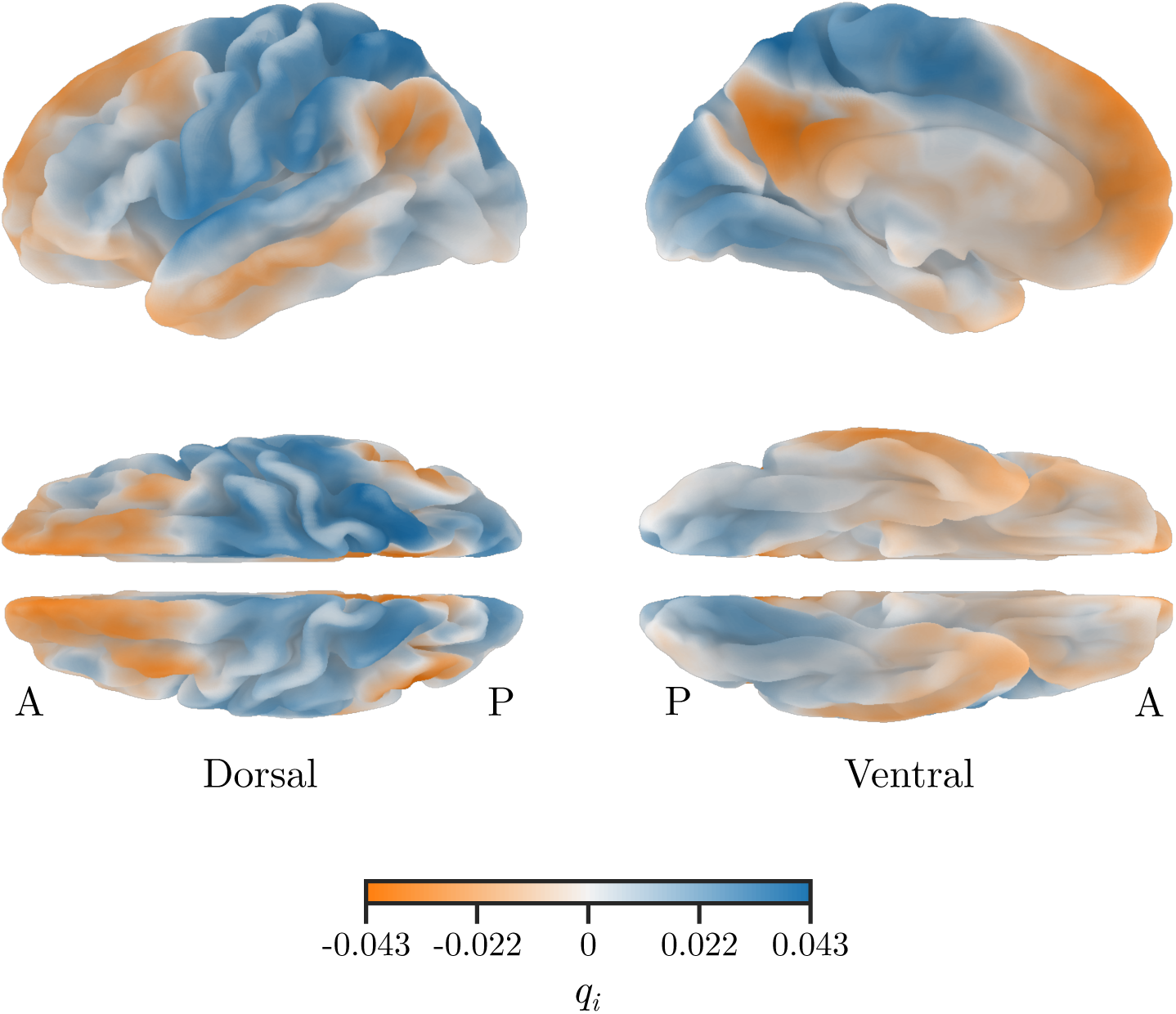
Brain surface map of the MMD partition gradient for the Diez dataset projected onto the Freesurfer pial surface template. Color indicates the interpolated value of *q*_*i*_. A: anterior. P: posterior.

When the brain is faceted by lobe affiliation, several notable patterns emerge. The frontal lobe is demarcated into the prefrontal cortex (pfc) and pre-motor areas and dorsolateral pfc anatomically, which are respectively situated in opposite quadrants of the functional embedding. In addition, the parietal lobe is split largely into default mode network (dmn) regions – including regions of the inferior parietal lobule and precuneus – and primary and secondary unimodal areas, including somatosensory cortices and areas involved in visual processing. Consistent with this observation, the occipital lobe has the largest proportion of regions belonging to the putative tpn at 83.42%. Taken together, these data suggest that maximizing the mmd in the context of vectorized connectomes is able to recover biologically-relevant network characteristics, while also accounting for the presence of negative edges, thereby removing heuristic steps that may bias downstream analyses as a result.

## 4 Discussion

In this study we presented a novel graph embedding approach for rs-fmri connectivity using rest2vec. Rest2vec improves upon current methods by using the full range of correlative information and representing the functional relationships of the brain in a low-dimensional embedding. Whereas many processing strategies involve arbitrary thresholds, rest2vec does not involve removing any data from the functional connectome. Previous studies have suggested that these negative correlations may have important – but still not fully understood – biological roles [37]. While there exist variations of methods that account for negative edges, such as the Louvain algorithm [14] and the *Q*^∗^-maximization method [9], the issue of deciding the appropriate weight of contribution to assign to these edges still remains.

Previous work from our group demonstrated that using a probability-based divisive approach with permutation testing could recover the hierarchical community structure of rs-fmri connectomes while preserving negative edges, which we called probability-associated community estimation (pace) [10]. In addition, our previous study [4] demonstrated how nonlinear dimensionality reduction and manifold learning techniques could be used to investigate the intrinsic geometry of structural connectomes derived from diffusion imaging. Inspired by these approaches, we sought to develop a method by which rs-fmri functional connectomes could be represented in their intrinsic geometry while also preserving negative edge relationships.

Dimensionality reduction techniques have been previously applied to neuroimaging datasets, e.g., clustering in lower dimensions to demarcate subjects belonging to different clinical populations, such as healthy controls and patients. Here, rest2vec applies dimensionality reduction at the level of brain regions. Furthermore, we chose to use the isomap method because it uses a geodesic distance metric for generating the lower-dimensional embedding [8]. By doing so, distance in the lower-dimensional embedding conveys meaningful information, as opposed to other methods, such as *t*-sne [38], that are stochastic and primarily meant for clustering purposes.

In the context of functional connectivity, converting the coordinate system to a polar representation was an intuitive visualization decision, as it centers the data around the origin where regions with lower Θ_*i,j*_ values are mapped closer to the origin and regions with higher Θ_*i,j*_ values are mapped in the periphery. Interestingly, regions with a greater number of high Θ_*i,j*_ values (i.e., more out-of-phase relationships) tended to be unimodal and also have low within-cluster Θ_*i,j*_ values, as seen most clearly in the occipital lobe (SI Figure 15). In contrast, more centrally-embedded regions tended to be located in brainstem regions (known to facilitate various sensory relay roles) and associative regions. This is reminiscent of Mesulam’s synaptic hierarchy model [39], where primary unimodal regions are embedded at the periphery, most proximal to sensory input, with downstream synaptic connectivity progressing inward towards the center to heteromodal and associative areas.

By using lower-dimensional embedding distance metrics, we were able to recover functionally relevant relationships. In the case of the occipital lobe, mapping the intrinsic functional distance to its cluster centroid in the isomap embedding generated a gradient map in the anatomical space of the dorsal and ventral visual streams [33, 34]. On the dorsal surface, the gradient proximal to the occipital lobe can be seen going to the posterior parietal regions, whereas on the ventral surface the proximal gradient extends from the occipital lobe to the inferior temporal lobe (Figure 5). In another example, the precuneus had two primary clusters in the isomap embedding. When projected onto the brain surface, these two clusters demarcated the dorsal-anterior and ventral-posterior portions of the precuneus (Figure 7). The dorsal-anterior gradient appeared to primarily consist of the superior parietal, somatomotor, and occipital cortices. The ventral-posterior gradient appeared to be composed of the posterior cingulate, parahippocampal, and superior occipital cortices and the hippocampus. There is evidence for the dorsal-anterior and ventral-posterior portions of the precuneus being involved in different functions. A rs-fmri study by [36] identified the dorsal and anterior portions of precuneus having stronger connectivity with areas including the occipital, somatomotor, and posterior parietal cortices and the superior temporal gyri. In addition, they identified the ventral precuneus as being more strongly associated with the middle frontal gyrus, posterior cingulate cortex, cuneus, and calcarine sulcus. This demarcation is thought to be due to the diverse roles of the precuneus. In particular, the dorsal-anterior portion of the precuneus, which has strong connectivity with the occipital and superior parietal cortices, is involved in processing polymodal imagery and visuospatial information, whereas the ventral-posterior precuneus is thought to be more involved in episodic memory retrieval [35]. While the study by [36] further subdivided the precuneus into eight clusters in their study, our results were largely consistent with their observations, suggesting that rest2vec can detect heterogeneous connectivity patterns within individual regions.

In addition to representing the intrinsic geometry of functional connectomes, we proposed using the maximum mean discrepancy (mmd) method by [13] to partition the connectome into maximally functionally distinct modules. The mmd was originally implemented to detect how different two probability distributions were to test if they were from the same population [13]. For our use case, we maximized the mmd as an objective function to find two populations of brain regions such that their distributions are as distant as possible to identify functional communities. One advantage is this is a vectorized approach and does not rely on iterative methods. In addition, this method offers flexibility in the choice of probability and kernel similarity measures used as input, and so are not limited to only Pearson correlation measures.

When the functional connectome is represented in its intrinsic embedding using nonlinear dimensionality reduction, the mmd partition elicited a strikingly symmetric representation. Upon closer observation, these two communities were split approximately between the canonical task-positive network (tpn) and the default mode network (dmn), consisting of the precuneus, inferior parietal lobule (ipl), posterior cingulate cortex, hippocampus, and areas of the prefrontal cortex (pfc), among others [40]. This initial bifurcation is consistent with previous modularity studies [10], and is a validation that this embedding procedure is capturing functionally-relevant characteristics. In addition, we showed lobe-specific affiliations for the two communities. These results were consistent with the putative dmn/tpn split. Notably, the ipl and precuneus are shown in contrast to the postcentral regions within the parietal lobe; similarly, the pfc and pre-motor areas show clear boundaries. Together, these results demonstrated that using this mmd approach to solve the connectome modularity problem yielded reproducible and biologically-meaningful connectome partitions, and that the properties of these communities can be represented using dimensionality reduction.

### Limitations and future directions

In this paper, we used rs-fmri connectomes from a group of subjects in order to compute the probability of there being a negative correlation between each pairwise edge between regions. While this approach led to consistent results across two independent datasets, we did not assess how robust this procedure was to intersubject variability or the size of groups. In addition, while average rs-fmri connectomes yield a wealth of functional connectivity information, they are a static representation of a dynamic process. Furthermore, there has been increasing emphasis on individual connectome analysis with aims towards personalized medicine [41, 42]. To that end, future improvements on these methods will need to incorporate dynamic as well as subject-specific analyses of functional connectivity. Recent works by [43, 44] have suggested the concept of hierarchical or multi-scale networks, which could lead to natural extensions of this work via subject embedding spaces which are in turn composed of network embedding spaces.

Another limitation is we only examined the mmd partition at the first bifurcation into two communities. While this proved effective as a proof-of-concept, further work will need to be done to develop a hierarchical way to detect *N* communities with this approach. In addition, more robust methods could be used for maximizing the mmd objective function to avoid the possibility of local maxima to achieve better accuracy.

### Conclusion

Rest2vec incorporates both positive and negative edge connectivity using a model inspired by statistical mechanics to transform functional connectome data into phase angle relationships. This representation of the connectome can be combined with nonlinear dimensionality reduction techniques to represent the intrinsic geometry of the functional connectome in a lower-dimensional embedding. Together, these methods allow for a vectorized approach to investigate the functional relationships of rs-fmri brain connectivity data. In addition, we connected rest2vec to the maximum mean discrepancy metric to demonstrate how rest2vec can be used to address the modularity problem as a kernel two-sample test. In summary, we presented a rs-fmri connectome graph embedding technique that uses nonlinear dimensionality reduction and statistical learning methods to create a low-dimensional representation of the intrinsic geometry of the functional connectome.

## Acknowledgements

This study was supported by NIH AG057468 to ZDM, NIH AG056782 to LZ and ADL. The authors declare no conflicts of interest.

## 5 Appendix: Supplemental information

**Figure 13:**
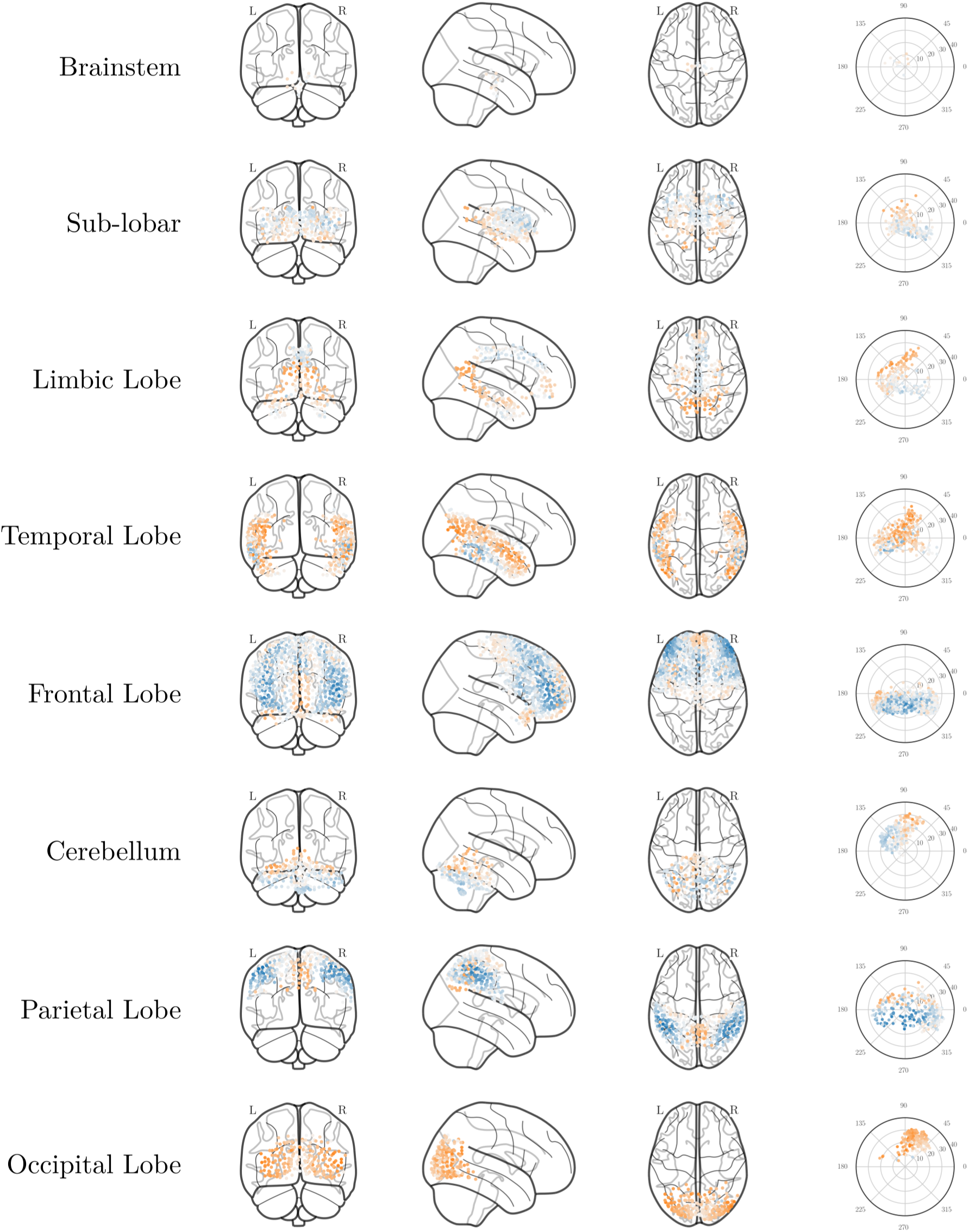
Anatomical and functional embedding of the Diez dataset faceted by anatomical lobe affiliation. Color indicates the partition predicted by the eigenvector of 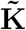 corresponding to the second-highest eigenvalue.

**Figure 14:**
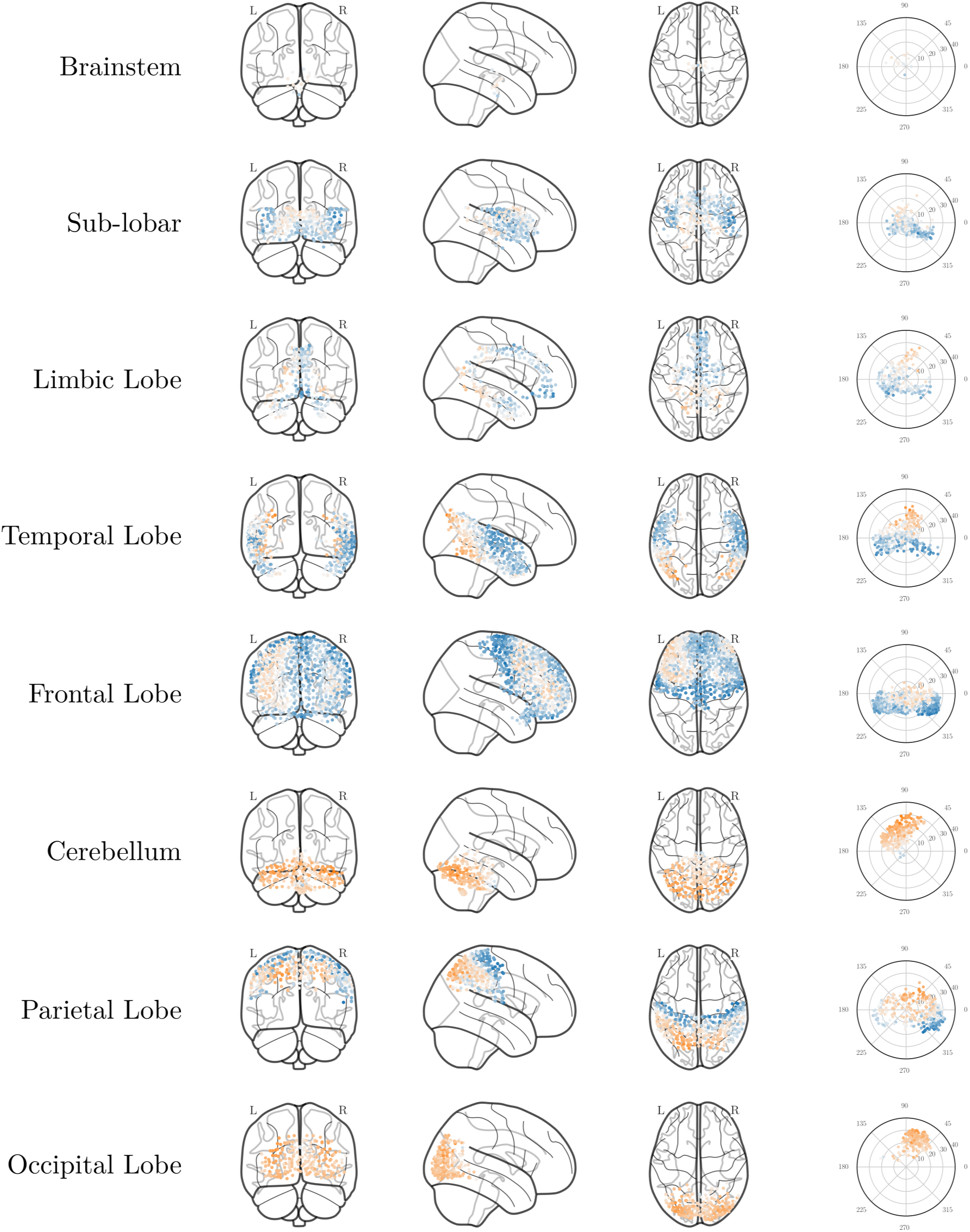
Anatomical and functional embedding of the Diez dataset faceted by anatomical lobe affiliation. Color indicates the partition predicted by the eigenvector of 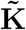 corresponding to the third-highest eigenvalue.

**Figure 15:**
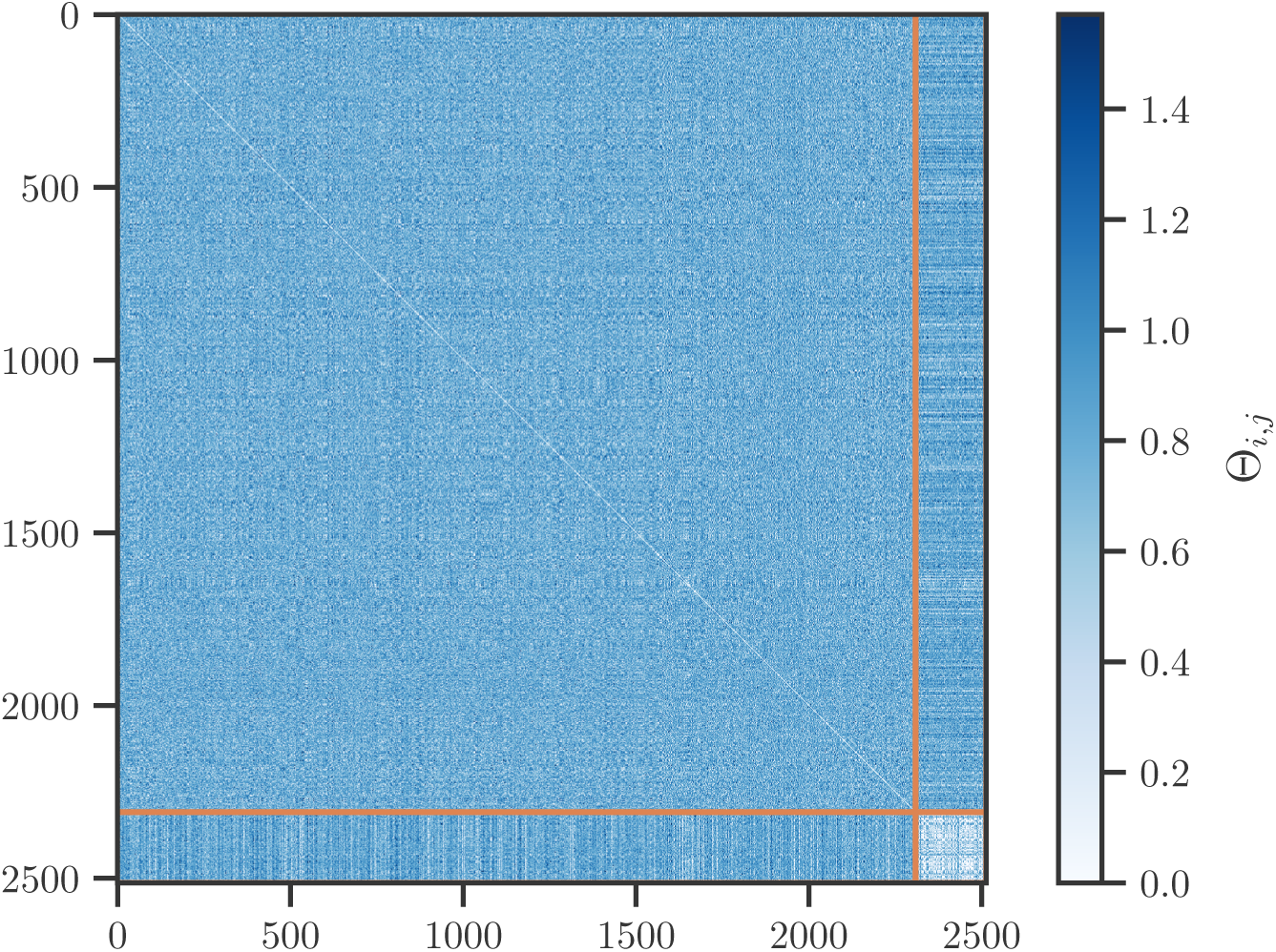
Θ for the Diez dataset with rows and columns arranged in a diagonal grid into non-occipital lobe regions (top left block) and occipital lobe regions (bottom right). Boundary between blocks is indicated by the orange line.

**Figure 16:**
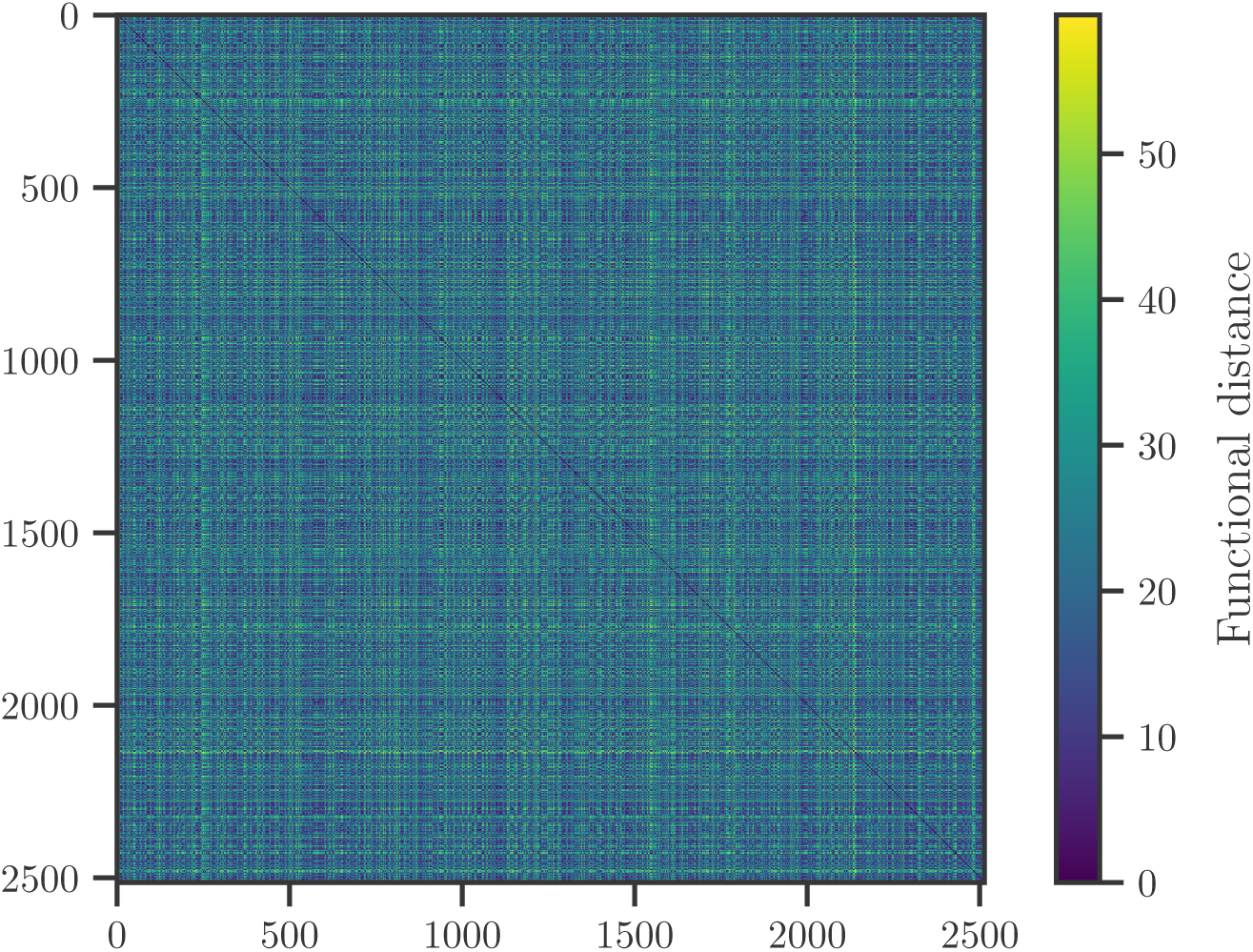
Pairwise functional distance for the Diez dataset.

